# In Utero CFTR Modulation Alleviates Disease in G551D Cystic Fibrosis Pigs

**DOI:** 10.64898/2026.01.20.698888

**Authors:** Sarah E. Ernst, David K. Meyerholz, Melissa S. Samuel, Kristin M. Whitworth, Youssef W. Naguib, David S. Nakhla, Mahmoud H. Abou Alaiwa, Christoph O. Randak, Qian Dong, Lynda S. Ostedgaard, Tayyab Rehman, Brie M. Hilkin, Linda S. Powers, Mal R. Stroik, Nicholas D. Gansemer, Michael R. Rector, Peter J. Taft, Rachel Hedinger, Brian J. Goodell, Stephen E. Mather, Ramkrishna Sen, Ian M. Thornell, Steven A. Bullard, Raissa F. Cecil, Josh A. Benne, Jamison J. Ash, Linda D. Boyken, Phillip H. Karp, Ping Tan, Shu Wu, Anthony J. Fischer, Ashley L. Cooney, Patrick L. Sinn, Alejandro A. Pezzulo, Kiho Lee, Paul B. McCray, Joseph Zabner, Aliasger K. Salem, Randy S. Prather, Michael J. Welsh, David A. Stoltz

**Affiliations:** Departments of Internal Medicine, Roy J. and Lucille A. Carver College of Medicine University of Iowa, Iowa City, Iowa 52242; Departments of Pathology, Roy J. and Lucille A. Carver College of Medicine University of Iowa, Iowa City, Iowa 52242; Departments of Pediatrics, Roy J. and Lucille A. Carver College of Medicine University of Iowa, Iowa City, Iowa 52242; Departments of Molecular Physiology and Biophysics, Roy J. and Lucille A. Carver College of Medicine University of Iowa, Iowa City, Iowa 52242; Pappajohn Biomedical Institute, Roy J. and Lucille A. Carver College of Medicine University of Iowa, Iowa City, Iowa 52242; Department of Biomedical Engineering, University of Iowa, Iowa City, Iowa 52242; Department of Pharmaceutical Sciences and Experimental Therapeutics, College of Pharmacy, University of Iowa, Iowa City, Iowa 52242; Howard Hughes Medical Institute, University of Iowa, Iowa City, Iowa 52242; Division of Animal Sciences, University of Missouri, Columbia, Missouri 65211; Department of Medicine, University of Michigan, Ann Arbor, MI 48109

## Abstract

Previous studies indicate that pigs with *CFTR-null* and *CFTR-ΔF508* mutations develop multiorgan disease similar to that in people with cystic fibrosis (CF). At birth, their airways exhibit host defense defects that predispose to airway infection, inflammation, and mucus accumulation. The *CFTR-G551D* mutation causes CF by producing CFTR channels that localize correctly but have reduced channel activity. Ivacaftor (VX-770) is a small molecule drug developed to potentiate CFTR activity. To test the phenotype of the *CFTR-G551D* mutation in pigs and determine whether ivacaftor can rescue CF abnormalities, we developed *CFTR^G551D/G551D^* (CF-G551D) pigs through homologous recombination in fetal fibroblasts and somatic cell nuclear transfer. Newborn CF-G551D piglets exhibited phenotypes similar to CF-null piglets, including meconium ileus, exocrine pancreatic destruction, micro-gallbladder, vas deferens destruction, and airway structural abnormalities. Compared to wild-type pigs, CF-G551D pigs had reduced forskolin-stimulated short-circuit current in airway and intestinal tissues. Ivacaftor increased the single-channel open state probability of *CFTR-G551D* and increased short-circuit current to near wild-type levels. Similar to our other CF pig models, we found that 100% of CF-G551D pigs were born with meconium ileus. To test whether *in utero* ivacaftor treatment could prevent or alleviate meconium ileus, pregnant sows were treated with ivacaftor beginning at day 35 of gestation and continuing until delivery. This treatment rescued the pancreas, gallbladder, and vas deferens phenotype in the majority of CF-G551D pigs. Animals that were spared from meconium ileus were able to survive without ivacaftor treatment. Airway disease developed similar to other CF pig models. These findings indicate that this model may be useful for studies in which CFTR function can be reversed, for investigating *in utero* CFTR correction strategies, and for longitudinal studies in CF pigs.

## INTRODUCTION

Cystic fibrosis (CF) is a life-threatening genetic disorder characterized by chronic respiratory infections, pancreatic insufficiency, and gastrointestinal disease, affecting 70,000 - 100,000 individuals worldwide (1). CF is caused by mutations in the *cystic fibrosis transmembrane conductance regulator* (*CFTR*) gene, which encodes an anion channel critical for maintaining electrolyte and fluid balance across epithelial surfaces (2–5). More than 2,000 CFTR genetic variants have been identified, with the *CFTR-G551D* mutation being one of the most prevalent gating mutations, accounting for approximately 4-5% of all CF cases (6, 7).

The *CFTR-G551D* mutation impairs the regulation of CFTR, leading to reduced chloride and bicarbonate secretion despite normal CFTR protein synthesis and trafficking (8, 9). This mechanism of dysfunction contrasts with the more common *CFTR-ΔF508* mutation, in which CFTR protein misfolding and degradation are predominant (10–12). Individuals with the *CFTR-G551D* mutation exhibit severe CF clinical manifestations, including progressive lung disease, gastrointestinal complications, and reduced life expectancy, underscoring the need for precise disease models (13, 14).

Animal models have significantly advanced the understanding of CF pathophysiology and therapeutic development. Murine models, while instrumental in elucidating basic CF mechanisms, fail to recapitulate key human CF features, such as spontaneous lung disease and pancreatic dysfunction, likely due to species-specific anatomical and physiological differences (15). Conversely, CF pigs exhibit a disease phenotype remarkably like humans, including meconium ileus, pancreatic insufficiency, and airway pathology, making them an ideal model for studying CF disease progression and therapeutic interventions (16, 17).

The development of a CF-G551D pig model is particularly compelling due to the mutation’s responsiveness to CFTR potentiators like ivacaftor (VX-770), which has transformed clinical outcomes for people with CF and the *CFTR-G551D* mutation (18–20). A pig model harboring the *CFTR-G551D* mutation would not only enhance our understanding of mutation-specific pathophysiology but also provide a valuable platform for preclinical testing of CFTR modulators and other emerging therapies.

Therefore, the objective of this study was to develop and characterize a CF-G551D pig model to more accurately recapitulate human CF disease and facilitate the investigation of genotype-specific therapeutic strategies. By bridging the translational gap between basic research and clinical application, this model holds the potential to (1) better understand the implications of *in utero* CFTR modulation, (2) further knowledge of CF disease pathogenesis, and (3) accelerate the discovery of effective treatments for individuals with CFTR gating mutations.

## METHODS

### Patch clamp experiments

Studies with porcine G551D CFTR were performed as previously described (21). Excised, inside-out membrane patches from 293T human embryonic kidney cells transiently expressing porcine pG551D CFTR using a pcDNA™3.1 (Invitrogen, Carlsbad, CA) CFTR expression system. The pipette (extracellular) solution contained: 140 mM N-methyl-D-glucamine, 3 mM MgCl_2_, 5 mM CaCl_2_, 100 mM L-aspartic acid, and 10 mM tricine, pH 7.3 with HCl. The bath (intracellular) solution contained 140 mM N-methyl-D-glucamine, 3 mM MgCl_2_, 1 mM Cs ethylene glycol bis (2-aminoethyl ether) – N,N,N’,N’ tetraacetic acid (CsEGTA), and 10 mM tricine, pH 7.3 with HCl. Following patch excision, CFTR channels were activated with 22 nM PKA catalytic subunit (from bovine heart, EMD Millipore Corporation, Billerica, MA) and ATP (magnesium salt, Sigma-Aldrich, St. Louis, MO). PKA catalytic subunit was present in all cytosolic solutions that contained ATP. Experiments were performed at room temperature (23 - 26°C). Recordings from patches containing 1-6 channels were digitized at 5 kHz and prior to analysis low-pass filtered at 500 Hz using an 8-pole Bessel filter (Model 900, Frequency Devices, Inc., Haverhill, MA). Single channel openings and closings were analyzed with a burst delimiter of 20 msec (22) using Clampfit software (version 10.3, Molecular Devices, Sunnyvale, CA). Events < 4 msec duration were ignored. For patches containing more than one channel the mean interburst interval (IBI) was calculated using the formula P_o_ = (BD x P_o,Burst_) / (BD + IBI), where P_o_ is the mean open state probability, BD is the mean burst duration and P_o,Burst_ is the mean open state probability within a burst (23).

### Targeting vector construction

A 5,500bp region of pig genomic DNA centered around CFTR exon 12 was amplified with Platinum^TM^ *Taq,* DNA Polymerase High Fidelity (Invitrogen), then cloned into the TOPO TA cloning vector (Invitrogen). The *CFTR-G551D* mutation was made using the QuickChange II site directed mutagenesis kit (Agilent). LoxP flanked reverse neomycin resistance cassette driven by a PGK1 promoter was inserted 74 bp downstream from exon 12 (legacy exon 11). To accommodate the packaging limits of recombinant adeno-associated virus (AAV), the homology arms of the construct were shortened to 1.2 kb upstream of the mutation and 1.2 kb downstream of the PGK1 promoter sequence. The total vector length was ∼4.5 kb. The resulting targeting vector sequence was inserted into the G0202 AAV shuttle plasmid provided by the University of Iowa Viral Vector core which contained AAV2 inverted terminal repeats. Plasmid DNA was purified, sequenced, and deposited at the Viral Vector Core at the University of Iowa for AAV2.1 virus production.

### Fetal fibroblast cell culture

Pig fetal fibroblasts were harvested from the skin of fetal piglets on day 35 of gestation. Tissue was digested for 12 hr in culture media composed of high glucose DMEM supplemented with 10% HyClone fetal bovine serum (FBS) (Cytiva), L-glutamine, non-essential amino acids, sodium pyruvate, and penicillin/streptomycin. For dissociation, 2 mg/ml collagenase (Sigma) was added to the culture media. Cells were passed through a 70 µM cell strainer (Corning) and washed once by centrifugation at 1000 x g with fresh culture media without collagenase. Cells were resuspended in freezing media composed of 90% FBS and 10% DMSO, then aliquoted into 1 mL cryovials. Cell freezing was controlled to -1°C/min using the Mr. Frosty^TM^ freezing container (ThermoFisher). The following day, cells were transferred to liquid nitrogen for later use.

### Fetal fibroblast infection and selection

1 x 10^6^ fetal cells were cultured in a 100 mM biotin coated tissue culture dish with fetal fibroblast media. Cells were infected with 5 x 10^11^ particles of rAAV2.1. After 12 hr the cells were detached using TrypLE Express (Gibco) dissociation media. Dissociated cells were divided into fifty 96-well plates. Divided cells were allowed to reach confluency (48 hr) and then underwent antibiotic selection with 100 µg/mL geneticin. After 10 d of antibiotic selection, cells were split. Half of the cell suspension was lysed for PCR screening, and the rest of the cell suspension was passaged into a new 96-well plate. Cells reserved for PCR screening were spun at 1,000 rpm in 96-well conical plates for 5 min. The supernatant was carefully aspirated to avoid dislodging the cell pellet, and cells were resuspended into 20 µl of lysis buffer. Cells were then lysed at 95°C for 10 min.

### Homologous recombination PCR screen

Cells were screened in 96-well format using Platinum^TM^ *Taq* DNA Polymerase (Invitrogen), and PCR primers designed with the forward primer located in intron 11 outside of the vector homology arm and the reverse primer located in the neo cassette. PCR fragments from screened positive cells were submitted for sequencing to ensure the *CFTR-G551D* mutation was present. G551D positive cells were further screened to ensure the downstream sequence was correct using a second primer outside of the vector homology.

### Somatic cell nuclear transfer and herd generation

Cells containing the *CFTR-G551D* mutation were frozen and sent to the University of Missouri where they were cloned by somatic cell nuclear transfer into wild-type surrogate sows as described previously (24, 25). Genetic material was isolated from piglet tails and each piglet was genotyped by PCR using primers designed outside the homology arms of the targeting vector. PCR products were then sequenced to ensure the presence of an unmodified allele as well as a mutated allele. Resulting piglets were heterozygous for G551D. Heterozygous animals were mated resulting in *CFTR^+/+^*; *CFTR^+/G551D^*; and *CFTR^G551D/G551D^*piglets. Additionally, several litters were made by breeding a *CFTR^+/G551D^* sow and a *CFTR^G551D/G551D^*boar who was maintained on ivacaftor treatment throughout life (including *in utero* treatment).

### *In utero* treatment with ivacaftor

Ear fibroblasts from *CFTR-G551D* homozygous fetal piglets euthanized at 90 days gestation were cultured and cloned as previously described (24–26) and injected into wildtype surrogate sows. The sows were fed 5 mg/kg ivacaftor twice daily starting at day 35 gestation until full term C-section delivery or allowed to deliver naturally. Ivacaftor (Matrix Scientific) was complexed with (2-hydroxypropyl)-β-cyclodextrin (Sigma-Aldrich) at a ratio of 1:2 ivacaftor:cyclodextrin (27) then suspended in yogurt. After the initially cloned animals displayed intestinal phenotype correction, heterozygous animals were mated to produce natural litters resulting in homozygous G551D animals exposed to ivacaftor *in utero*.

### Ussing chamber studies

Trachea, bronchus and nasal turbinate tissues were cultured as previously described (24–26) and analyzed in modified Ussing chambers (Physiologic Instruments) to determine ion transport properties. For studies done with symmetrical chloride conditions, cultures were bathed in bilateral high chloride buffer containing 135 mM NaCl, 5 mM HEPES, 2.4 mM K_2_HPO_4_, 0.6 mM KH_2_PO_4_, 1.2 mM CaCl_2_, 1.2 mM MgCl_2_, and 5mM dextrose, and bubbled continuously with compressed air. Chemical interventions included addition of 100 µM amiloride (Sigma), 100 µM DIDS (Sigma), 10 µM forskolin (Cayman Chemical), 10 µM ivacaftor (Selleck Chemicals), 100 µM Gly H-101, and 100 µM bumetanide (Sigma). All drugs were dissolved in DMSO and diluted 1:1000 in the apical chamber, except for bumetanide which was added to the basolateral side.

### Intestinal electrophysiology

Ileal segments from a newborn animal were excised immediately after euthanasia and mounted into Ussing chambers bathed bilaterally with a Krebs bicarbonate ringers solution containing 118.9 mM NaCl, 25 mM NaHCO_3_, 2.4 mM K_2_HPO_4_, 0.6 mM KH_2_PO_4_, 1.2 mM CaCl_2_, and 1.2 mM MgCl_2_. Initially, 5 mM dextrose was added on the basolateral side only, and tissues were bubbled continuously with a 5% CO_2_, 21% O_2_, nitrogen balanced gas mixture. Chemical intervention included 10/100 µM forskolin/IBMX (DMSO/ethanol), and 10 µM ivacaftor (DMSO). All drugs were added apically.

### Ivacaftor dose response

Cultured tracheal cells from a person with CF (genotype: *CFTR*^Δ*F508/G551D*^) and a *CFTR^G551D/G551D^* piglet were mounted into Ussing chambers. The apical surface was bathed in a low chloride solution containing 135 mM sodium gluconate, 2.4 mM K_2_HPO_4_, 0.6 mM KH_2_PO_4_, 1.2 mM CaCl_2_, 1.2 mM MgCl_2_, and 5 mM dextrose on both sides. The high chloride solution described for previous culture studies was added to the basolateral side. Cultures were bubbled continuously with compressed air. Chemical intervention included the addition of 100 µM amiloride, 100 µM DIDS, 10 µM forskolin, and increasing doses of ivacaftor starting from 1 nM up to 100 µM. Data was normalized to 100% for the final dose in cultures from each species.

### RNA isolation and qPCR measurements

Tissue samples were dissected and immediately placed into RNALater (ThermoFisher) and flash frozen with liquid nitrogen. Samples were stored at -80°C until RNA isolation could be performed. RNA was isolated with the RNeasy lipid and tissue kit (Qiagen). Sample quality was verified by the University of Iowa Genomics Institute. RNA concentration was measured with the Nanodrop spectrophotometer, and 2000 ng of RNA was reverse transcribed using Vilo Superscript reverse transcription kit (Invitrogen). cDNA was then diluted 1:20 using 10 ng of cDNA per reaction. qPCR primers were designed to amplify CFTR regions exon 11-13. β-actin was used as the housekeeping gene. Experiments were performed on the Quant Studio 6 Flex (Applied Biosystems) with the Fast SYBR^TM^ green qPCR Master Mix (Applied Biosystems).

### Ivacaftor pharmacokinetic and dose finding experiments

A piglet weighing approximately 10 kg was given 20 mg/kg of ivacaftor complexed with (2-hydroxypropyl)-β-cyclodextrin (Sigma-Aldrich) at a ratio of 1:2 ivacaftor:cyclodextrin (27). Ivacaftor was suspended in milk replacer and delivered by mouth (PO). Blood samples were collected at various time points post drug administration up to 24 hr. Plasma was isolated and ivacaftor levels were measured for each collection. Pregnant sows weighing approximately 150 kg were given series of four doses of 5 mg/kg fed every 12 hours mixed with high fat yogurt. Blood samples were collected at various time points for 2 d. Sows were euthanized 42 hr after the initial dose. Fetal blood collection was not possible, so fetal liver was collected and homogenized for fetal ivacaftor measurements.

### Sample preparation for LC-MS/MS ivacaftor measures of plasma or liver homogenates

Approximately 1 ml of whole blood was collected from young piglets (several ml for adult animals) using sodium heparin blood collection tubes. Multiple aliquots of whole blood were centrifuged at 2000 x g to separate plasma from blood cells. The upper plasma layer was carefully transferred to a clean Eppendorf tube and frozen at -80°C for batch processing. Previously frozen plasma was thawed on ice and 20 µL of plasma was aliquoted into a new tube. 80 µL of acetonitrile (Sigma) was added to each tube, then vortexed for 30 sec to precipitate plasma proteins. Samples were centrifuged at 10,000 x g at 4°C for 20 min. 10 µL of the cleared supernatant was transferred to a clean Eppendorf tube, carefully avoiding disturbance of the protein pellet. 90 µL of pure water (Invitrogen) was added to each tube. After vortexing completely, the resulting 100 µL solution was transferred to glass sample tubes compatible with LC-MS/MS autosampler. Fetal liver samples were thawed on ice and weighed. Tissues were diluted 1:4 (w/v) in ultrapure water and homogenized until uniform. 20 µl of tissue homogenate was processed using the protocol described for plasma samples. All samples were submitted to the High-Resolution Mass Spectrometry Facility at the University of Iowa for analysis.

### ASL pH measurement

To measure ASL pH of cultured airway epithelia, a ratiometric pH indicator, SNARF-1, conjugated to 70 kD dextran (Thermo Fisher Scientific) was used. SNARF-1-dextran was delivered as a powder to the apical side and allowed to distribute into ASL for 1 hr. Imaging was performed on a laser-scanning confocal microscope Zeiss LSM 880. The microscope chamber housing epithelia maintained a humidified environment with 5% CO_2_ at 37°C. SNARF-1 was excited at 514 nm, and emissions at 580 nm and 640 nm were recorded. Five different SNARF-1 containing ASL regions were randomly selected per culture. Fluorescence emission ratios (580/640) were recorded, converted into pH using standard curves, and averaged to obtain ASL pH value per culture.

### Bacterial-coated grid killing assay

Bacterial killing assay was performed as previously described (28). Briefly, bacteria-coated grids were placed on the apical surface of primary cultures of differentiated airway epithelia. After exposure, grids were immediately rinsed with PBS and then immersed in PBS containing the fluorescent indicators SYTO 9 and propidium iodide (Live/Dead BacLight Bacterial Viability assay, Invitrogen). After 15 min, the grids were rinsed with PBS and placed on slides for imaging with a laser-scanning confocal microscope (Olympus FV1000).

### Histopathological analyses

At necropsy, tissue samples were harvested and immersed in 10% neutral buffered formalin for approximately seven days. Subsequently, the tissue specimens underwent dehydration processing using sequential alcohol and xylene baths before being embedded in paraffin. The embedded samples were cut into sections of approximately 4 μm thickness and mounted on glass Plus slides. Sections were then stained using two methods: hematoxylin and eosin (HE), and Periodic acid Schiff following diastase pretreatment (dPAS).

### Statistical Analysis

Data were analyzed using GraphPad Prism 10. *P* < 0.05 was considered statistically significant.

## RESULTS

### Ivacaftor restores CFTR function in porcine G551D

Excised, inside-out membrane patches from 293T human embryonic kidney cells transiently expressing porcine G551D-CFTR were used to test channel activity and ivacaftor responsiveness. As expected, porcine CFTR with the *G551D* mutation showed very little channel activity in the presence of PKA and ATP alone. However, activity significantly increased with the addition of 10 µM ivacaftor (**Figure 1A**). Similar to earlier human CFTR studies (**Figure 1B**), the open channel probability (P_o_), the fraction of time a channel is in the open state, increased significantly, the average time channels remained open also increased, and time between channel opening was dramatically reduced (**Figure 1C**). Together these data suggested that ivacaftor was likely to potentiate porcine G551D-CFTR function *in vivo*.

**Figure 1.**
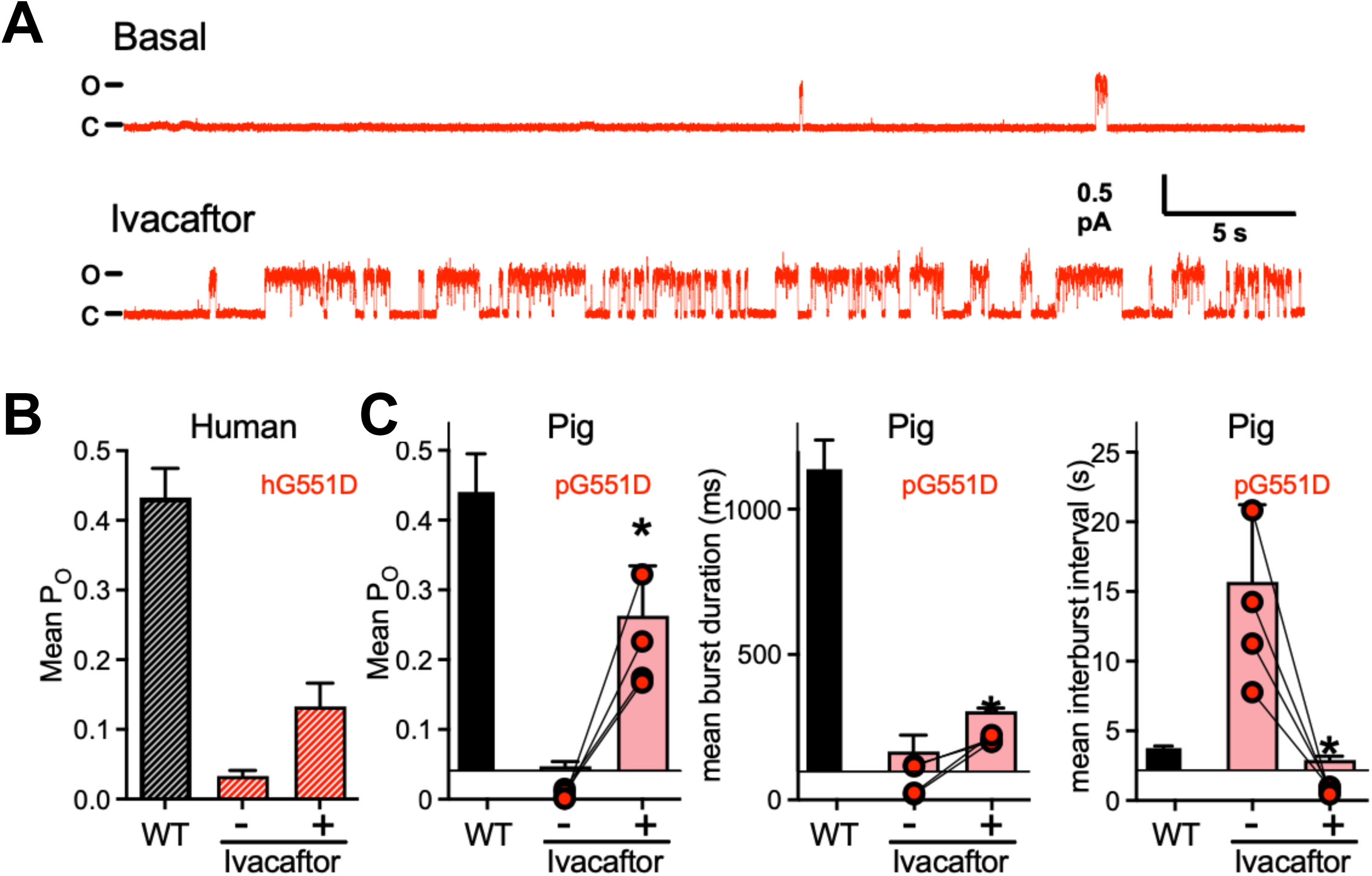
Influence of Ivacaftor on Single-Channel Gating of Porcine CFTR-G551D. Data are all from single-channel recording of excised inside-out membrane patches treated with 1 mM ATP and PKA. Data in red are from patches with CFTR-G551D, and data in black are from patches with wild-type CFTR. (**A**) Recording of pig CFTR-G551D before and after addition of 10 μM ivacaftor to the cytosolic surface of the patch. Holding voltage was -70 mV. For illustration purposes, traces were digitally low-pass filtered at 100 Hz. c, closed state; o, open state. P_o_ indicates open state probability. (**B**) Human wild-type and G551D CFTR (data are adapted from Van Goor *et al.* PNAS 2009). (**C**) Single-channel properties of porcine wild-type and G551D CFTR. (Wild-type data are adapted from Ostedgaard *et al.* PNAS 2007). Data for CFTR-G551D are results from 4 individual membrane patches. Bars are means with standard deviation. Asterisks indicate p < 0.05 compared to - ivacaftor.

### Development and phenotype of CF-G551D pigs

Fibroblasts from four 35-day-old fetuses were infected with recombinant AAV2.1 containing 2.4 kb of pig genomic DNA sequence homologous to the region around CFTR exon 12 containing the *G551D* mutation plus a neomycin resistance cassette. After infection, ∼30% of colonies proved to be antibiotic resistant. PCR screening was performed on fifty 96-well plates for each set of fibroblasts. The first two rounds of fibroblast screening did not identify any *G551D* positive clones. The final two screens identified a total of 18 *G551D* positive colonies. These cell lines were used for somatic cell nuclear transfer (SCNT). The first two attempts did not result in successful pregnancy, but the subsequent three attempts resulted in one male and two female cloned litters of *CFTR^+/G551D^* piglets (**Supplemental Figure 1**).

*CFTR^+/G551D^* piglets were mated when sexually mature and piglets homozygous for the *CFTR-G551D* mutation were evaluated soon after birth. Like previous models of CF in pigs (17, 24, 26), all CF-G551D newborn piglets had lethal meconium ileus, intestinal atresia/micro-colon, exocrine pancreatic destruction, micro-gallbladder, absent vas deferens, and tracheal structural abnormalities (**Figure 2**).

**Figure 2.**
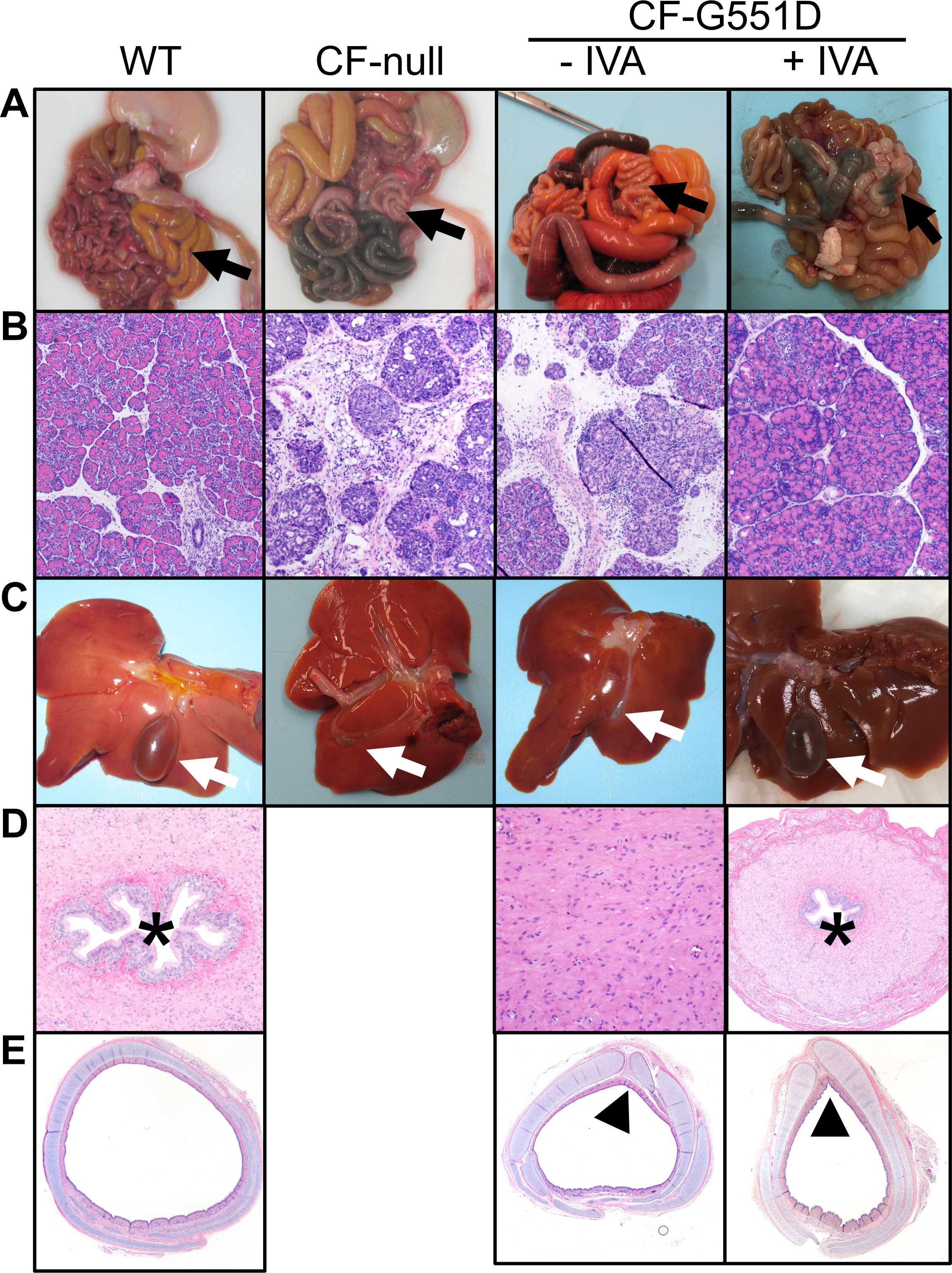
CF-G551D Pigs Develop CF Disease and *In Utero* Ivacaftor Treatment Prevents Disease Development. Gross and histological images. **(A)** Gastrointestinal tract. **(B)** Pancreas. (**C**) Gallbladder. (**D**) Vas deferens. (**E**) Trachea. Black arrows denote spiral colon. White arrows denote gallbladder. Asterisks denote vas deferens lumen. Black arrowheads denote cartilage ring defects.

### *CFTR* expression is similar between wild-type (WT) and CF-G551D tissues

*CFTR* mRNA levels were evaluated by qPCR in trachea, bronchus, and intestinal tissues from WT and CF-G551D piglets. In all tissues studied, *CFTR* expression was similar between WT and CF-G551D samples (**Figure 3A**). CFTR immunostaining in airway epithelial cultures revealed comparable amounts of CFTR protein that localized to the apical membrane (**Figure 3B**).

**Figure 3.**
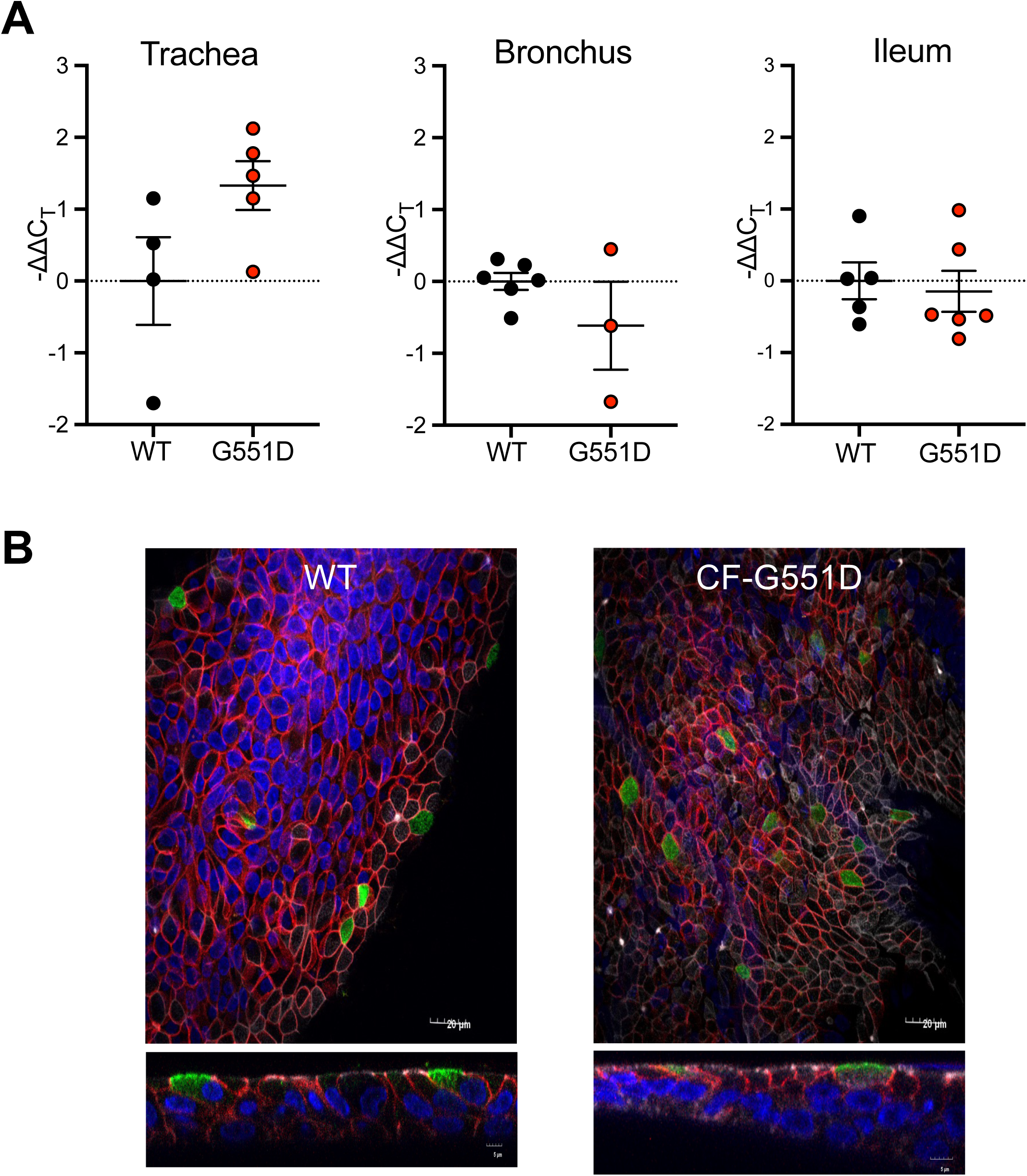
CFTR Expression is Similar in WT and CF-G551D Pigs. (**A**) *CFTR* mRNA from excised trachea, bronchus and ileum. Each symbol represents a sample from an individual animal. (**B**) CFTR immunostaining in cultured airway epithelia. CFTR (green), β-catenin (red), and nuclear stain DAPI (blue). *En face* (upper) and cross-sectional (lower) images displayed.

### Ivacaftor potentiates CFTR-mediated anion transport in CF-G551D epithelia

Patch clamp experiments with ivacaftor showed increased CFTR activity in single G551D-CFTR channels (**Figure 1**). We tested G551D-CFTR function in cultured airway epithelia and freshly excised tissues from WT and CF-G551D pigs (**Figure 4A**). In the absence of ivacaftor, forskolin increased short-circuit current in CF-G551D airway epithelial cultures, but the response compared to controls was significantly reduced and varied by tissue type (**Figure 4B-D**). In tracheal and turbinate cultures, ivacaftor increased short-circuit current to near WT levels, whereas in ethmoid cultures the response was ∼40% of WT. Forskolin induced changes in transepithelial conductance were limited (∼25% of WT) in tracheal and turbinate cultures. Ivacaftor rescued conductance to ∼60% of WT in tracheal cultures and ∼70% in nasal turbinate cultures. Ethmoid sinus cultures showed a very limited conductance to forskolin, but this response was enhanced with ivacaftor (**Figure 4D**). Ivacaftor sensitive conductance was reduced in ethmoid cultures compared to epithelia from trachea and turbinate. Similar to CF-G551D pig airway cultures, ivacaftor potentiated CFTR-mediated anion transport in excised intestinal tissue samples to near WT levels (**Figure 5**). Finally, to determine if ivacaftor had a similar potency for human and porcine CFTR-G551D, we compared the ivacaftor dose response in cultured airway epithelia from CF-G551D piglets and human ΔF508/G551D cultures. Ivacaftor’s potency was similar in human and pig epithelia. The EC_50_ was 3.1 x 10^-7^ M in human epithelia versus 3.6 x 10^-7^ M in pig epithelia (**Figure 6**).

**Figure 4.**
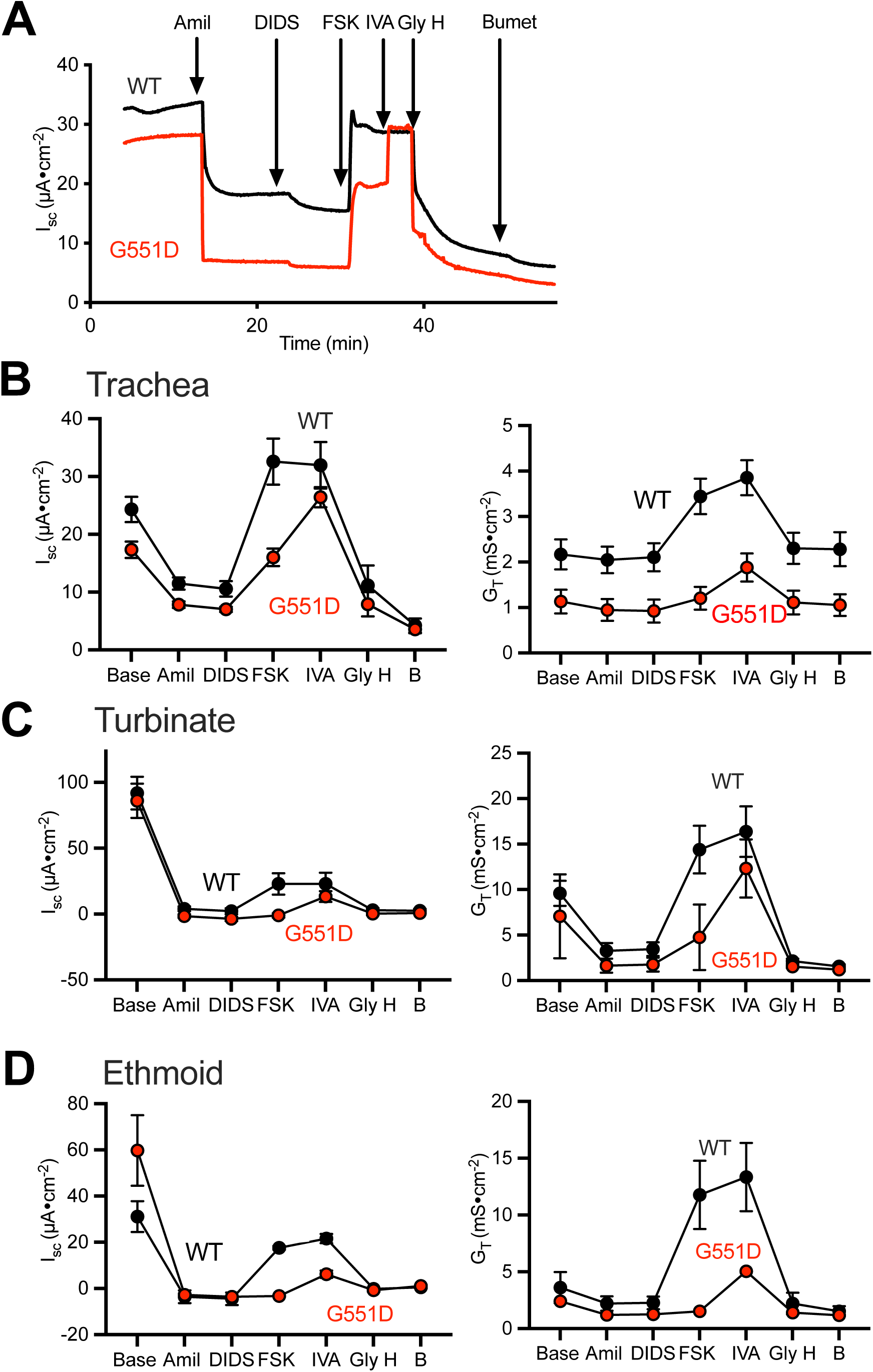
Ivacaftor Potentiates Porcine CFTR-G551D Airway Epithelia. (**A**) Representative current traces in response to indicated agents in WT and CF-G551D airway epithelia. (**B-D**) Short-circuit current (I_sc_) and conductance (G_T_) measurements from WT (black) and G551D (red) trachea, turbinate, and ethmoid airway epithelial cultures. Studies were performed with symmetrical chloride conditions with the addition of 100 µM amiloride (Amil), 100 µM DIDS, 10 µM forskolin (FSK), 10 µM ivacaftor (IVA), 100 µM GlyH1-01 (GlyH), and 100 µM bumetanide (B).

**Figure 5.**
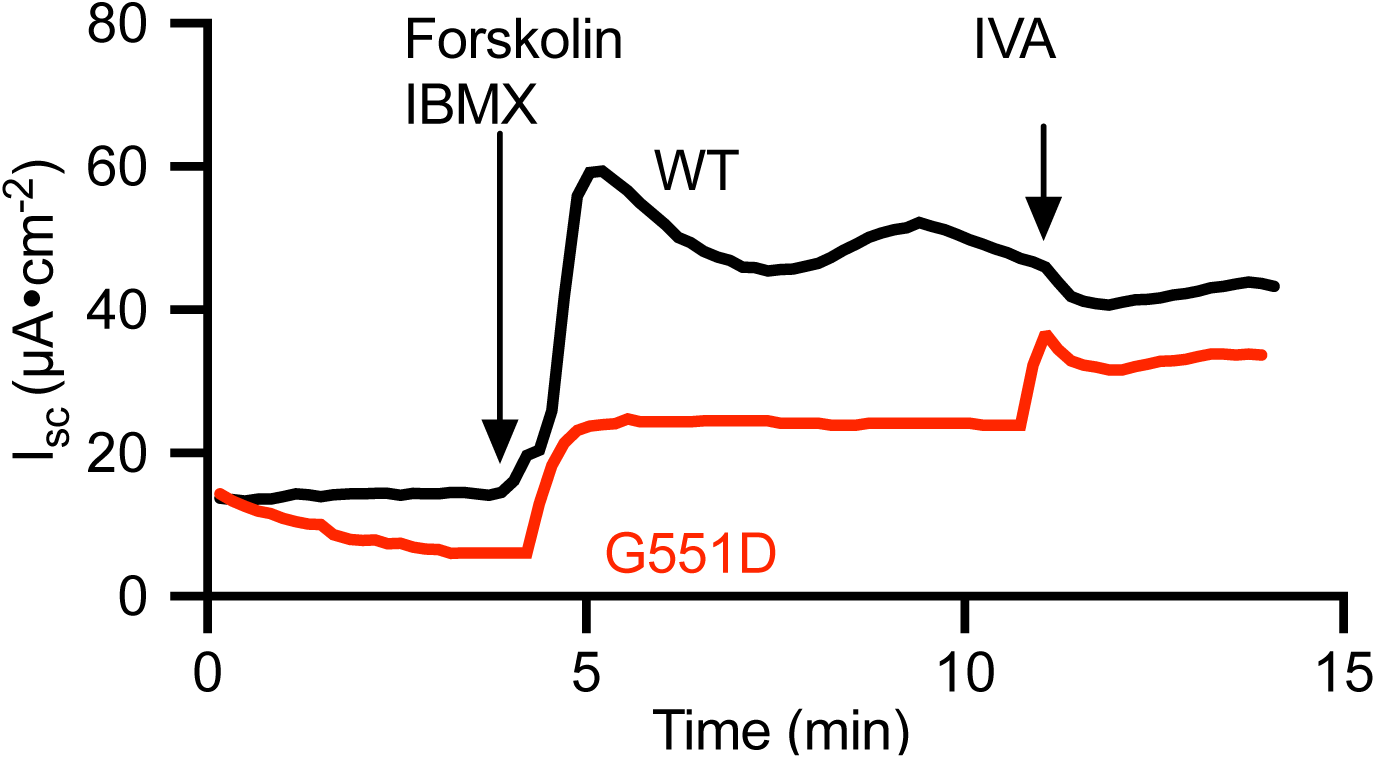
Ivacaftor Potentiates Intestinal CFTR-G551D. Electrophysiology of excised piglet intestine measured in Ussing chambers. Excised segments WT (black) and G551D (red) of ileal tissues were mounted in modified Ussing chambers and studied under symmetrical chloride conditions buffered with bicarbonate. Tissues were exposed to 10 µM forskolin (FSK) and 100 µM IBMX and 10 µM ivacaftor (IVA). A representative tracing is shown.

**Figure 6.**
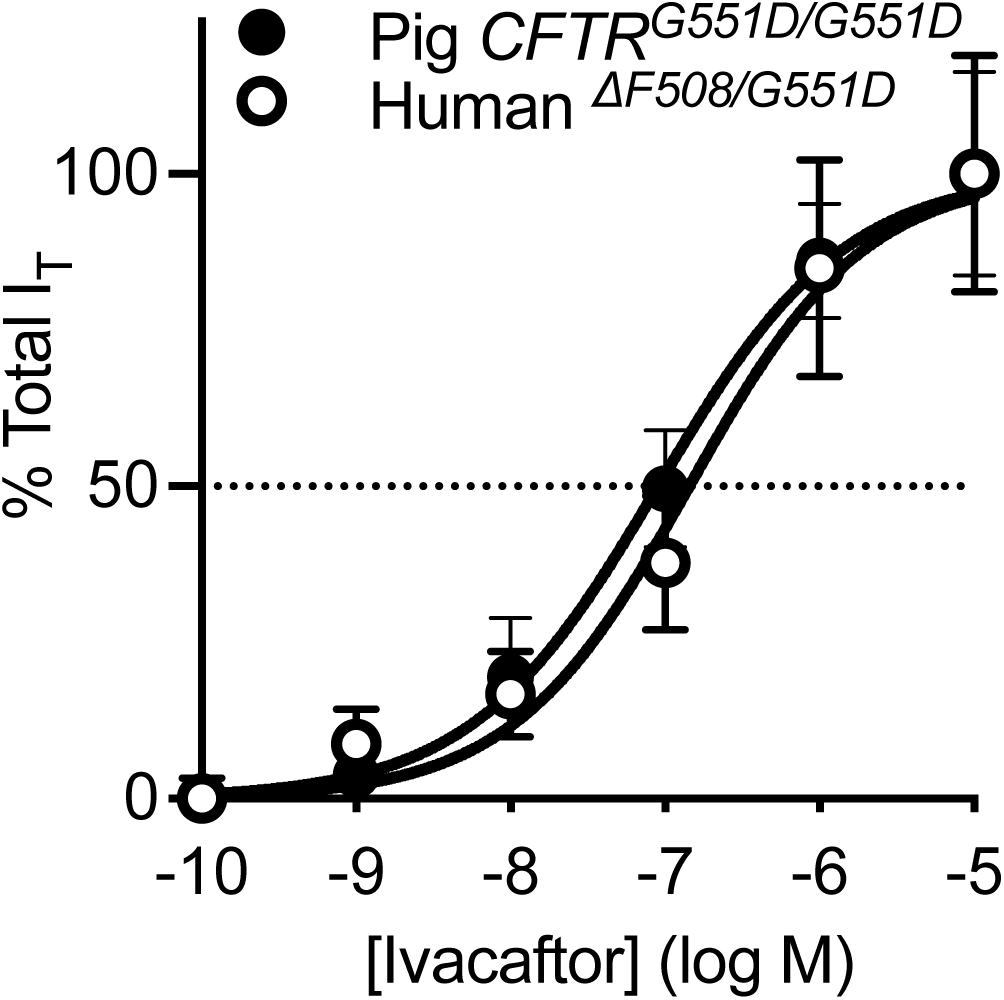
Ivacaftor has a Similar Potency for Human and Porcine CFTR-G551D. Ivacaftor dose-response in cultured trachea and bronchus from a CF human and CF-G551D piglets. Increasing doses of ivacaftor were added apically in the presence of amiloride (100 µM) and forskolin (10 µM) with low apical chloride conditions.

### Ivacaftor rescues CF host defense defects in cultured airway epithelia

We previously reported that neonatal CF pigs have a more acidic airway surface liquid (ASL) pH compared to non-CF (28). This more acidic ASL pH is linked to reduced ASL-mediated bacterial killing and mucociliary transport. We hypothesized that CF-G551D airway epithelial cultures would have a more acidic ASL pH and impaired bacterial killing and that ivacaftor could reverse these defects. We tested the effect of ivacaftor on ASL pH in cultured epithelia using a fluorescent pH indicator, SNARF (28). In the presence of forskolin, the ASL pH in CF-G551D epithelial cultures was reduced compared to WT levels (**Figure 7A**). Ivacaftor addition to CF-G551D cultures increased ASL pH. Bacterial killing was impaired in CF-G551D airway epithelial cultures; ivacaftor restored killing to WT levels (**Figure 7B**).

**Figure 7.**
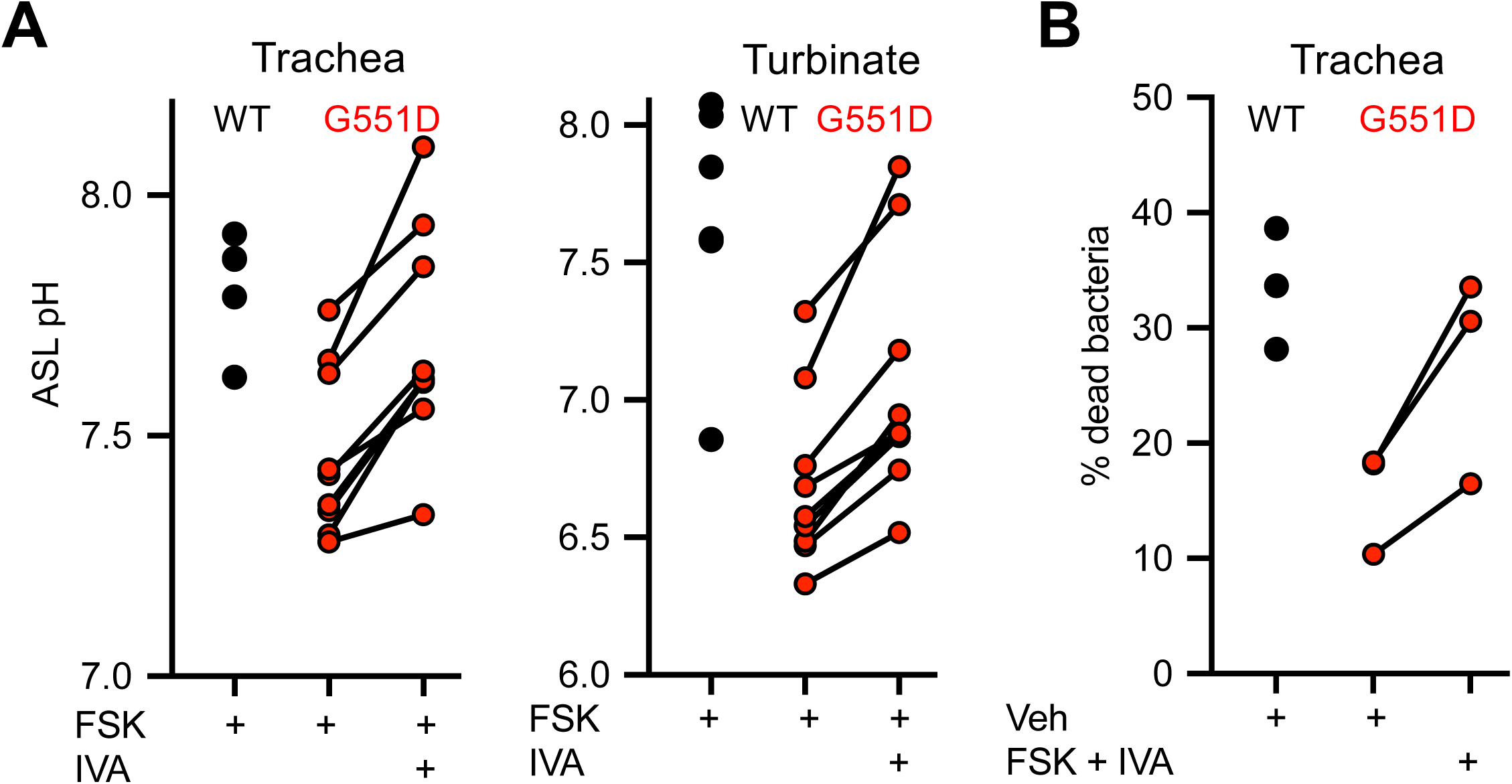
Ivacaftor Increases Airway Surface Liquid pH and Enhances Bacterial Killing in CF-G551D Airway Epithelia. (**A**) ASL pH was measured in WT (black) and G551D (red) airway epithelia with 10 µM forskolin (FSK) in the absence or presence of 10 µM ivacaftor (IVA). (**B**) WT (black) and G551D (red) airway epithelial cultures were exposed to *S. aureus* on gold grids in the presence or absence of forskolin and ivacaftor. Each symbol represents an epithelial culture from a different animal.

### *In utero* ivacaftor treatment alleviates CF disease in CF-G551D pigs

These data suggested that *in utero* ivacaftor treatment could rescue CFTR-G551D function *in vivo* and alleviate and/or prevent the lethal meconium ileus phenotype. We tested this prediction by treating pregnant sows carrying CF-G551D clones with ivacaftor. To determine the optimal ivacaftor dosing formulation and regimen, we first administered ivacaftor (20 mg/kg) orally to ∼ 10 kg control pigs. Since ivacaftor is highly lipophilic and nearly insoluble in water, ivacaftor was complexed with cyclodextrin (27). Ivacaftor plasma concentration levels peaked within 2 hr at 3.2 µM and remained detectable 24 hr later (**Supplemental Figure 2A**). This level is similar to earlier studies in humans. We also fed ivacaftor (5 mg/kg, every 12 hr) to a WT pregnant sow (∼150 kg) to determine the pharmacokinetics. Sow plasma ivacaftor levels rose over time (**Supplemental Figure 2B**). After 4 doses of the oral formulation (48 hr), we harvested fetal liver tissue. Liver homogenate ivacaftor concentrations were ∼3.0 µM (**Supplemental Figure 2B**). This level is comparable although slightly less than 4.12 µM reported in people taking ivacaftor.

A WT surrogate sow carrying CF-G551D clones was treated with 5 mg/kg ivacaftor, twice daily, starting around 35 days gestation (pig gestation period/pregnancy is ∼114 days) and continuing to term. Plasma levels of ivacaftor were monitored at several points throughout gestation to ensure continued drug absorption. The litter was born via cesarean delivery and produced 3 live piglets and 1 stillborn piglet. One piglet was euthanized shortly after birth to evaluate the gastrointestinal phenotype. Necropsy revealed significantly improved intestinal phenotype (**Figure 2A**) and a relatively normal appearing pancreas (**Figure 2B**), gallbladder (**Figure 2C**), and vas deferens (**Figure 2D**), while trachea structural abnormalities persisted (**Figure 2E**). The two remaining piglets were able to pass meconium and survived until electively euthanized. We repeated this approach in another non-CF surrogate sow carrying CF-G551D clones and treated with ivacaftor. Two CF-G551D piglets were born in this litter with 1 having a favorable gastrointestinal phenotype (survived the neonatal period) while the other piglet required euthanasia due to intestinal blockage.

Once *CFTR^+/G551D^* pigs were sexually mature, we mated heterozygous animals resulting in *CFTR^+/+^*, *CFTR^+/G551D^*, and *CFTR^G551D/G551D^* piglets. In these litters, *in utero* ivacaftor treatment also alleviated the lethal meconium ileus phenotype but at a reduced rate compared to cloned litters (80% piglet survival in cloned litters versus 75% survival in heterozygous matings) (**Table 1**). From these matings, we kept a male CF-G551D pig alive and maintained him on ivacaftor treatment. At approximately 6 months of age, we collected ejaculate from this CF-G551D pig. The collected semen had viable sperm present (**Figure 8A**), confirming that both *in utero* and long-term post-natal ivacaftor treatment preserved the structure and function of the vas deferens/epididymis (**Figure 8B**). This CF-G551D boar was mated to several *CFTR^+/G551D^* heterozygote sows and the pregnant sows received *in utero* ivacaftor treatment. This mating resulted in *CFTR^+/G551D^*and *CFTR^G551D/G551D^* piglets. Although this mating strategy increased the number of CF-G551D piglets in each litter, we found that only 54% of CF-G551D piglets had a favorable intestinal phenotype. While we do not know the mechanism(s) for the reduced ivacaftor effect on the meconium ileus phenotype in the litters resulting from the *CFTR^+/G551D^*x *CFTR^G551D/G551D^* matings, we suspect that it is related to significant inbreeding. Therefore, in the future, it might be best for some studies to produce CF-G551D pigs via SCNT where we observed much higher rates of meconium ileus correction.

**Figure 8.**
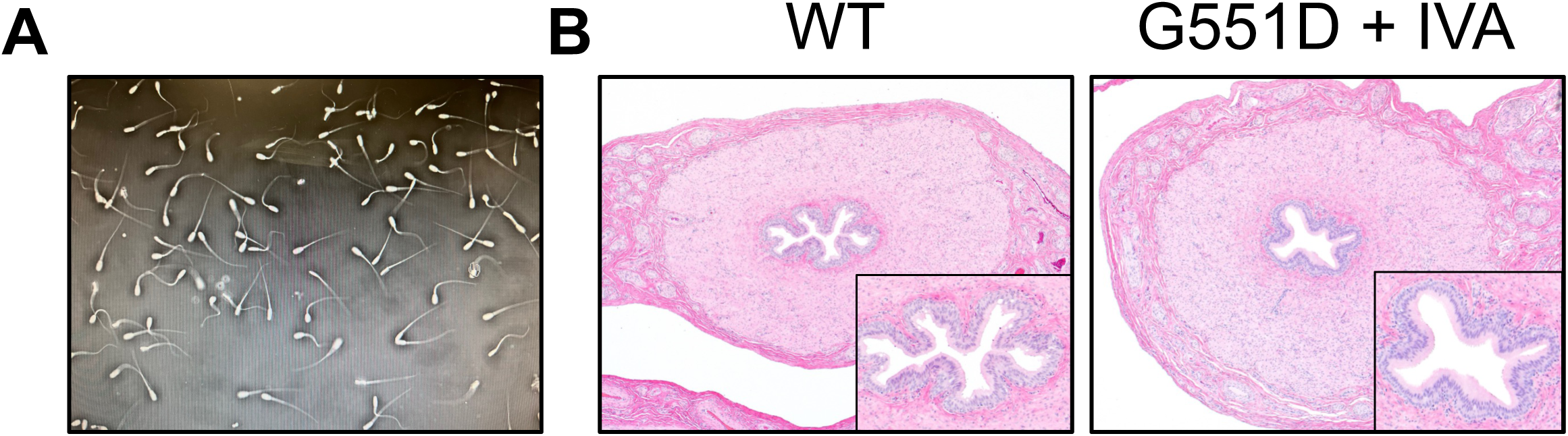
*In Utero* and Postnatal Ivacaftor Treatment Prevents Vas Deferens Destruction in Young CF-G551D Pigs. (**A**) Microscopic image of sperm in the ejaculate of a ∼6-month-old CF-G551D boar. (**B**) Histological images of the vas deferens from a WT and CF-G551D pig (5 months old) treated with ivacaftor *in utero* and postnatally.

**Table 1.**
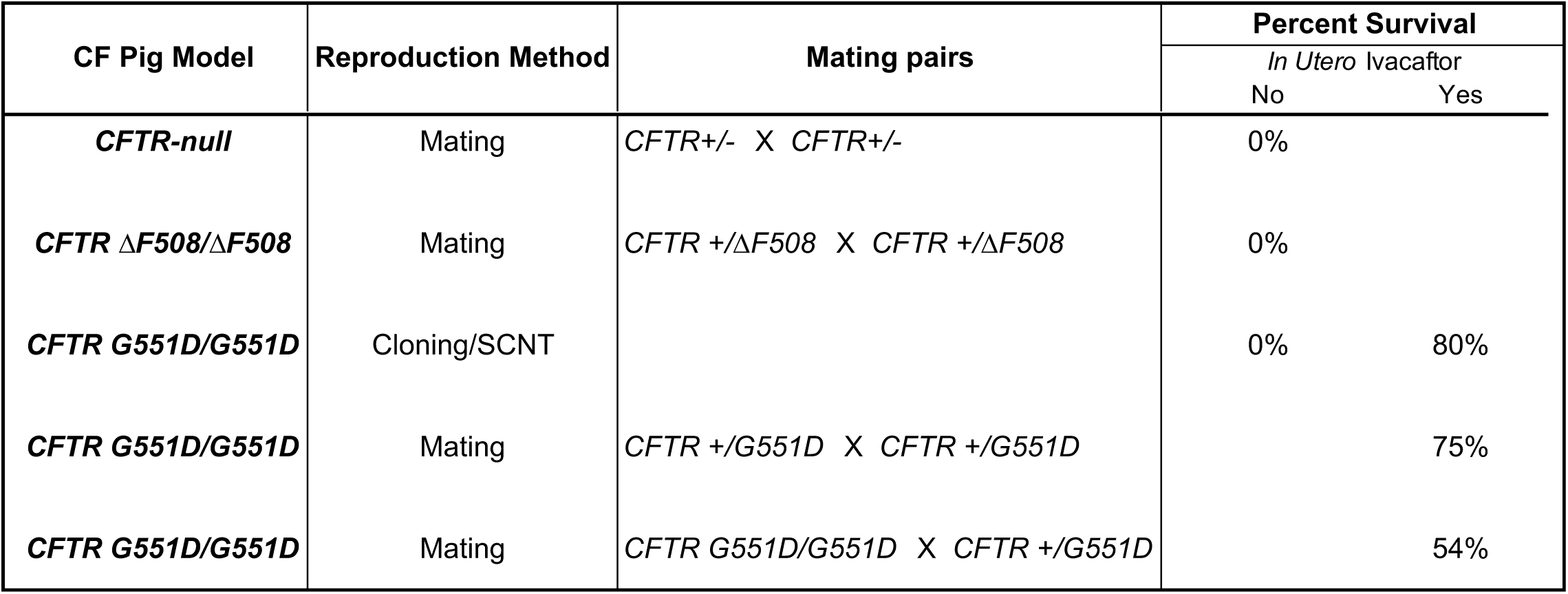

To determine if CF-G551D pigs develop lung disease similar to our other CF pig models, we studied some pigs that received no ivacaftor after birth. In these CF-G551D pigs we found evidence of CF lung disease including: (1) increased neutrophils, IL-1β, and bacteria (**Figure 9A-C**); (2) airway mucus and inflammatory cell accumulation (**Figure 9D-H**); and (3) radiographic evidence of air trapping and bronchial wall thickening (**Figure 9I-J**). Finally, we asked if continuation of ivacaftor postnatally would prevent disease development or lessen disease severity. Thus, we maintained some CF-G551D piglets on ivacaftor after birth. In a 2.5-month-old CF-G551D pig maintained on lifelong ivacaftor treatment, we saw minimal evidence of pancreatic disease (**Figure 10**).

**Figure 9.**
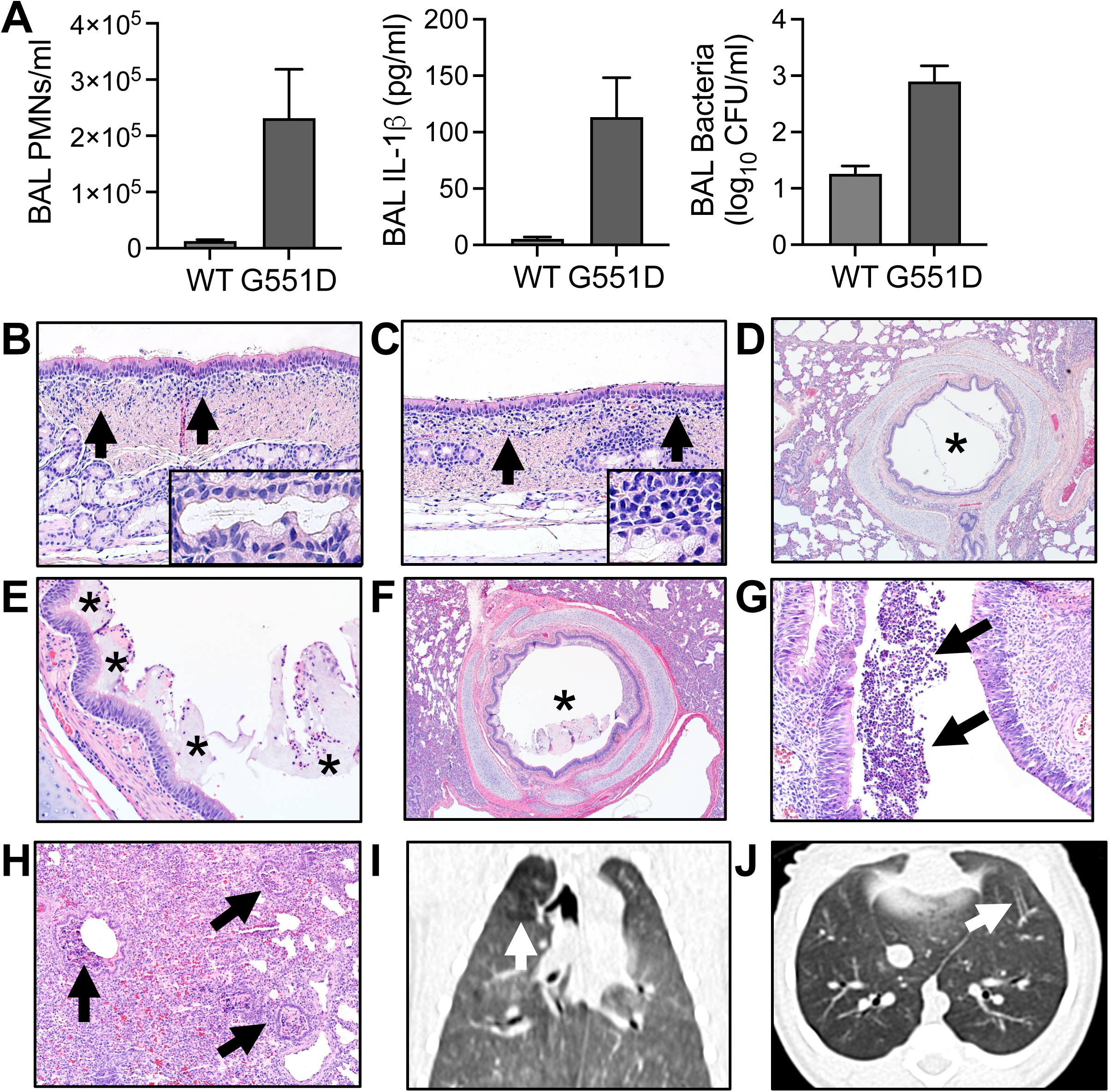
CF-G551D Pigs Develop Airway Disease in the Absence of Postnatal Ivacaftor Treatment. (**A**) Bronchoalveolar lavage fluid neutrophil (PMN) counts, IL-1β levels, and bacterial counts. n = three 10-day-old CF-G551D pigs. Mean ± SEM. (**B-H**) Histological images. **B** and **C** show airway epithelial inflammation (arrows) and submucosal gland duct dilation with ductal mucus accumulation. **D**, **E**, and **F** show luminal mucus and PMN accumulation (asterisks). **G** is an ethmoid sinus sample with accumulation of inflammatory cells. (**I-J**) Chest CT images. I shows air trapping (white arrow) and J shows airway wall thickening (white arrow).

**Figure 10.**
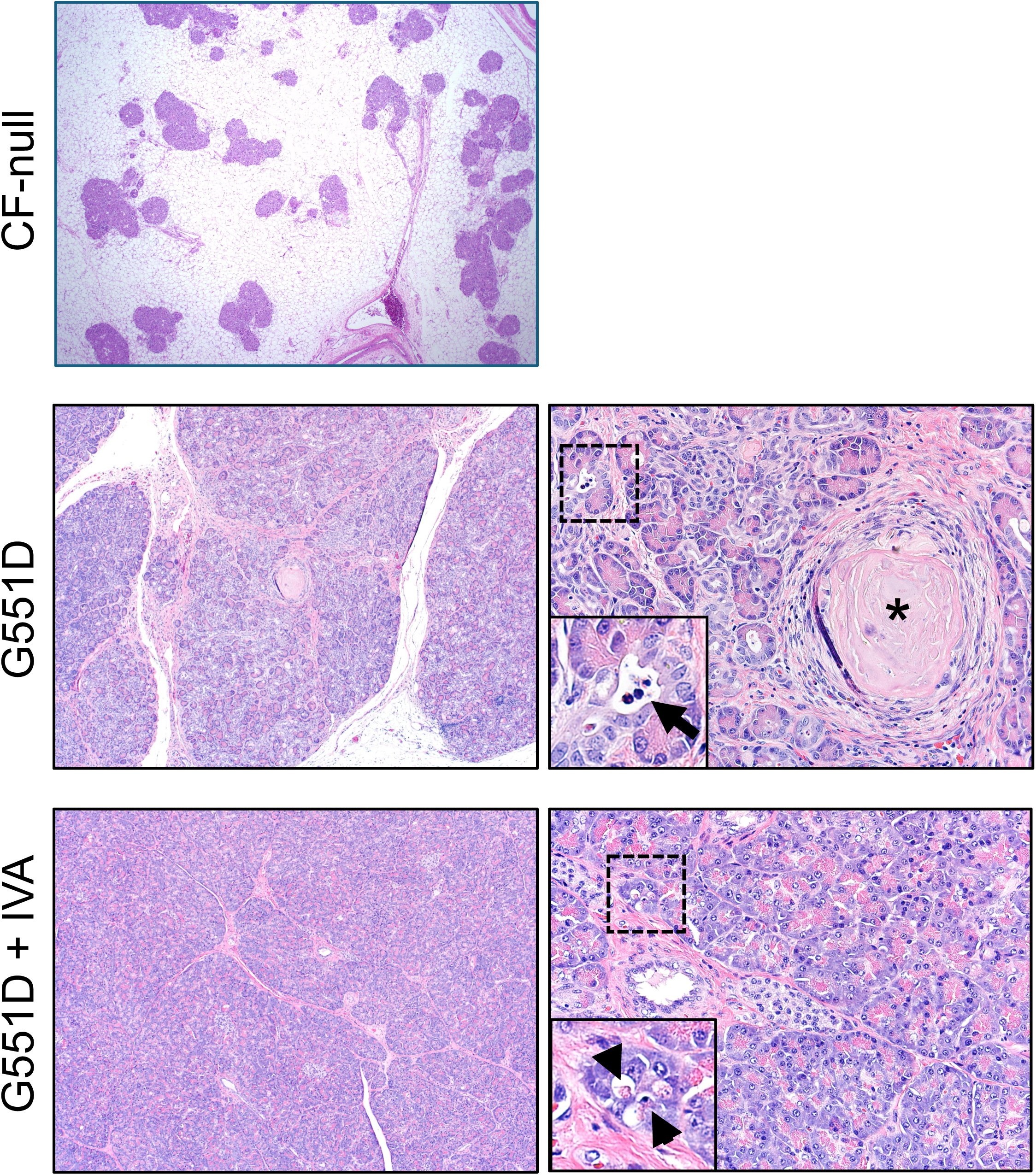
*In Utero* and Postnatal Ivacaftor Treatment Prevents Pancreas Disease in Young CF-G551D Pigs. Pancreas histological images from a ∼2-month-old CF-null pig and two 2.5-month-old CF-G551D pigs that received *in utero* ivacaftor treatment. In the CF-G551D pigs, postnatally ivacaftor treatment was either discontinued (G551D) or continued (G551D + IVA). CF-null pig pancreas was mostly composed of adipose tissue and little remnant pancreatic tissue. CF-G551D pancreas with *in utero* only treatment had mild to moderate active lesions. There was acinar/duct dilation and filling by eosinophilic proteinaceous obstruction material, scattered leukocytes (arrow, left inset) and cellular debris. In contrast, CF-G551D pancreas with continued ivacaftor treatment lacked lesions except for very rare vacuolar change in acini with some rare cellular debris (arrowheads, right inset) indicative of injury.

## DISCUSSION

We generated a porcine model of cystic fibrosis carrying the *CFTR-G551D* mutation. CF-G551D pigs exhibited hallmark features of human CF disease, including meconium ileus, exocrine pancreatic destruction, micro-gallbladder, vas deferens abnormalities, and tracheal structural defects. These findings demonstrate that the *CFTR-G551D* mutation produces a CF phenotype in pigs similar to that observed in *CFTR-null* and *CFTR-ΔF508* pigs. Importantly, *in utero* treatment with ivacaftor prevented or ameliorated many of these manifestations, including meconium ileus, pancreatic disease, and vas deferens abnormalities. Ivacaftor also restored defective electrolyte transport in CF-G551D tissues. These results provide further evidence that CFTR modulator therapy can prevent CF disease when initiated early in development.

The development of CFTR modulators has transformed the treatment of CF (19, 20, 29). However, questions remain about the optimal timing for initiating therapy and whether early intervention can prevent disease manifestations (30–32). The CF-G551D pig model provides a unique opportunity to address these questions. By administering ivacaftor during gestation, we demonstrated that correcting CFTR function *in utero* can prevent multiple CF manifestations. The prevention of meconium ileus is particularly notable, as this condition affects approximately 15-20% of infants with CF and is associated with increased morbidity (33, 34). Our findings suggest that early CFTR correction, before birth, may prevent disease in organs that undergo critical developmental processes during gestation.

The *CFTR-G551D* mutation is one of the most common gating mutations in CF, affecting approximately 4-5% of people with CF worldwide (6, 7). Unlike the *CFTR-ΔF508* mutation, which causes misfolding and degradation of CFTR, the *CFTR-G551D* mutation produces CFTR protein that reaches the cell surface but has severely impaired channel gating (8, 9). Ivacaftor was specifically developed to address this gating defect and was the first CFTR modulator approved for clinical use (19, 35, 36). By creating a pig model with the *CFTR-G551D* mutation, we generated a platform to test ivacaftor and potentially other potentiator compounds in a large animal model that closely mimics human CF disease. This is particularly important because the *CFTR-G551D* mutation responds well to ivacaftor in humans, and our pig model recapitulates this therapeutic response.

*In utero* CFTR modulator therapy represents an emerging area of CF research and treatment (37–42). In the CF-G551D ferret model, ivacaftor treatment initiated at embryonic day 28 (equivalent to the third trimester in humans) enhanced meconium passage from ∼20% in untreated animals to > 90% in treated offspring, while also preserving vas deferens development and partially protecting pancreatic function. Sustained postnatal ivacaftor treatment improved survival and slowed lung disease (39). Similarly, in the CF-ΔF508 ferret model, *in utero* and postnatal lumacaftor/ivacaftor treatment led to meconium passage in 100% of homozygous animals and provided partial protection from pancreatic insufficiency in 40% of treated ferrets, although treatment with ivacaftor alone was less effective (40). Recent case reports in humans and small case series have demonstrated that administration of elexacaftor/tezacaftor/ivacaftor (ETI) to pregnant carrier mothers can resolve fetal meconium ileus and may preserve pancreatic function in CF infants (37, 38, 41, 42). In a French multicenter study, 12 maternal-CF fetal dyads treated with ETI from the third trimester showed resolution of meconium ileus within a median of 14 days, though one fetus experienced increased bowel dilatation suggestive of intestinal atresia (41). Similarly, case reports from the United States and Europe have documented successful resolution of echogenic bowel and meconium ileus, along with false-negative newborn screening results and preserved pancreatic sufficiency in some infants exposed to ETI *in ute*ro (37, 38, 43). However, outcomes have been variable, with one Dutch case reporting persistent meconium ileus requiring surgery despite maternal ETI treatment, possibly related to timing of initiation, maternal body weight, or placental transfer variability (44). Safety data remain limited but reassuring, with no consistent pattern of major congenital anomalies reported, though isolated cases of cataracts, transient hypotonia, and neonatal pulmonary hemorrhage have been described. The French PROTECT workshop and ongoing studies such as MAYFLOWERS and MODUL-CF aim to systematically evaluate the safety, efficacy, and optimal dosing of *in utero* CFTR modulator therapy (38, 41, 45).

Our study revealed important insights about the timing and reversibility of CF disease. Some manifestations, such as meconium ileus and pancreatic disease, were largely prevented by *in utero* ivacaftor treatment. In contrast, we observed that certain abnormalities still developed despite treatment, including tracheal structural defects. These findings suggest that different organs have varying windows of susceptibility and that the timing of CFTR correction may be critical for some tissues. The pancreas appeared particularly responsive to early intervention, consistent with the idea that preventing early pancreatic damage might preserve endocrine and exocrine function (43, 46). Similarly, prevention of vas deferens abnormalities suggests that reproductive tract development requires functional CFTR during gestation. Understanding these developmental windows could inform clinical strategies for initiating modulator therapy in newborns and infants.

We do not know the optimal dose or timing for *in utero* CFTR modulator therapy. We initiated ivacaftor treatment once pregnancy was confirmed. Would initiating treatment at conception provide additional benefit? What is the minimum duration of treatment needed to prevent disease? Are there critical developmental windows during which CFTR function is most essential? The CF-G551D pig model provides a platform to address these questions. Future studies could investigate different treatment regimens, including varying doses, timing of initiation, and duration of therapy. Additionally, the model could be used to test combination therapies or next-generation modulators that may have improved potency or pharmacokinetic properties.

Our study has strengths and limitations. Strengths include: 1) We generated a porcine model with a human CFTR mutation that is clinically relevant and responsive to an approved therapy. 2) CF-G551D pigs exhibited disease manifestations similar to humans, including gastrointestinal, pancreatic, hepatobiliary, and reproductive abnormalities. 3) We demonstrated that ivacaftor treatment can prevent CF disease, providing validation of the model for therapeutic testing. 4) The large size and long lifespan of pigs allow for longitudinal studies and interventions that are not feasible in smaller animal models. 5) We were able to assess multiple organ systems and correlate therapeutic response across tissues. 6) The use of somatic cell nuclear transfer allowed us to generate pigs with a precise, defined mutation rather than relying on random mutagenesis.

Limitations include: 1) We only initiated ivacaftor treatment after pregnancy confirmation, which occurred several weeks after conception. Earlier treatment might have provided additional benefit or prevented abnormalities that persisted despite therapy. 2) The dose and pharmacokinetics of ivacaftor in pregnant sows and developing fetuses are not well characterized. We selected a dose based on previous studies and our preliminary work. 3) We did not investigate the long-term effects of *in utero* ivacaftor exposure, but no obvious defects were observed with our current approach. 4) The mechanisms by which ivacaftor prevents specific organ manifestations remain to be fully elucidated. 5) Only one CFTR mutation was studied, and findings may not generalize to other CF-causing mutations, particularly those that cause misfolding defects rather than gating defects.

In summary, CF-G551D pigs provide a unique opportunity to study a clinically relevant CFTR mutation and to test therapeutic interventions during fetal development. Our findings demonstrate that *in utero* ivacaftor treatment can prevent multiple CF manifestations, suggesting that early CFTR modulator therapy may have significant benefits. These results could have important implications for the timing of therapy initiation in newborns with CF and for the development of prenatal treatment strategies. The CF-G551D pig model represents a valuable platform for investigating CF pathogenesis, testing novel therapeutics, and optimizing treatment approaches to improve outcomes for people with CF.

