## Supplemental Figures for "In Utero CFTR Modulation Alleviates Disease in G551D Cystic Fibrosis Pigs"

### Supplemental Figure 1

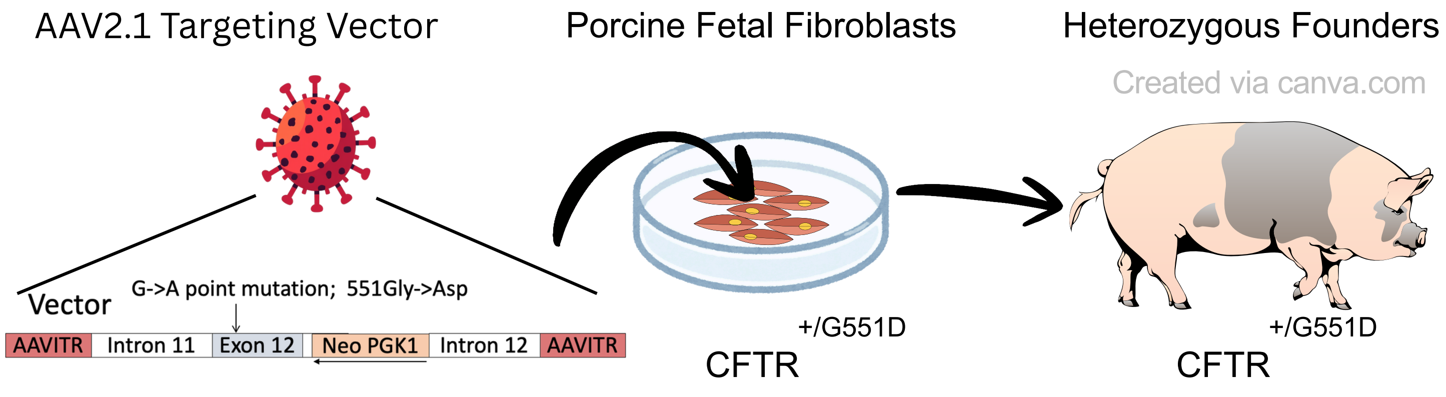


**Gene editing strategy via viral vector and homologous recombination.** Pig genomic DNA encompassing mutated CFTR exon 12 (G551D) was packaged into AAV2.1. Porcine fetal fibroblasts were infected with the viral vector and then screened for appropriate sequence integration. Genetic DNA from fibroblasts containing the G551D mutation was used to generate pigs heterozygous for the G551D mutation.

### Supplemental Figure 2

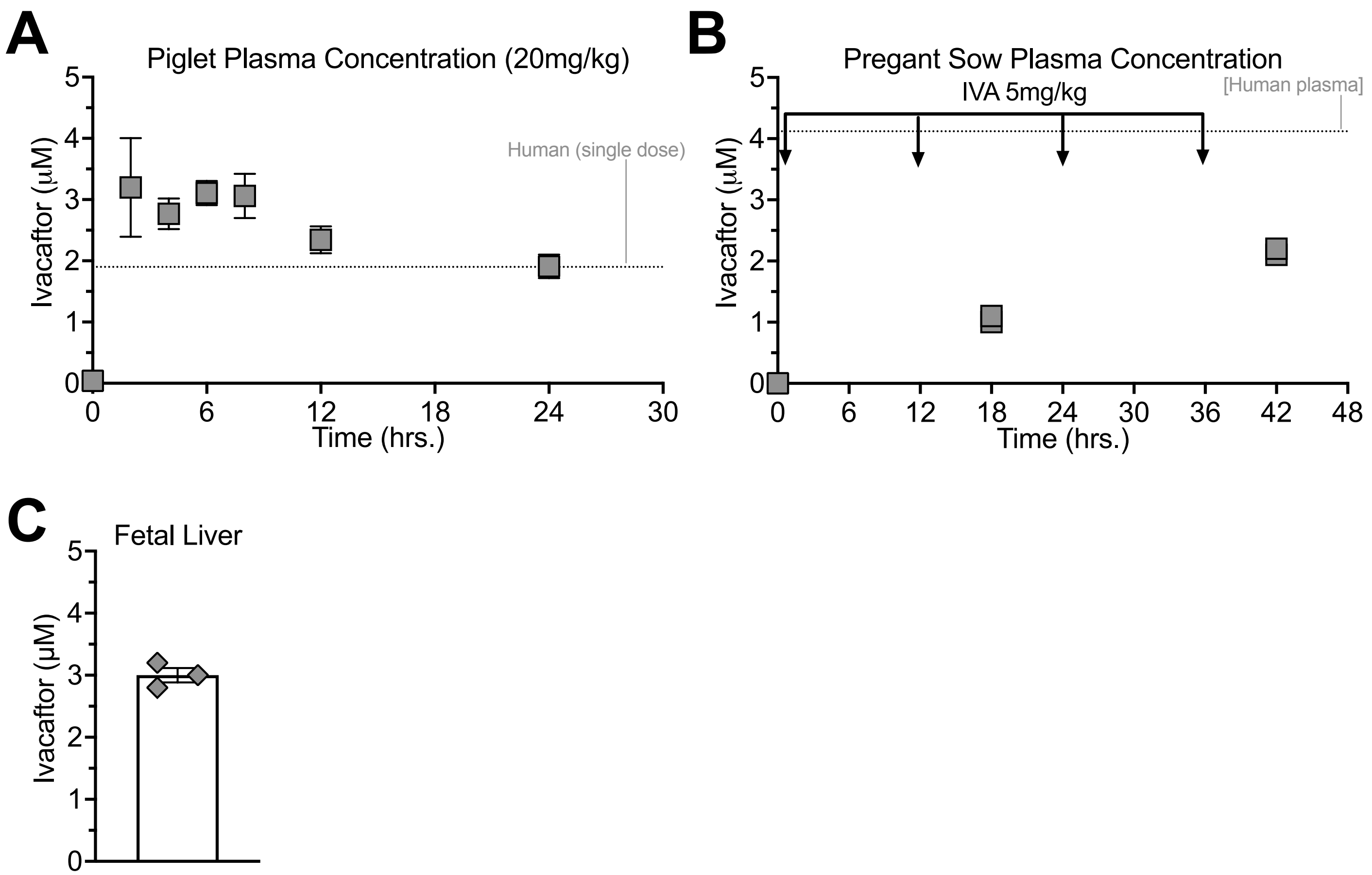


**Plasma ivacaftor concentrations in treated piglets, pregnant sows and fetuses are comparable to those reported in people with CF** (**A**) Wild-type piglets weighing approximately 10 kg were fed a single dose of ivacaftor complex (20 mg/kg). Blood samples were collected just before drug administration and at 2, 4, 6, 8, 12 and 24 hr post ivacaftor. Triplicate samples were measured after acetonitrile extraction by LC-MS/MS. Average values +/- SEM are displayed. (**B**) A pregnant sow was fed ivacaftor (5 mg/kg) given every 12 hr over two days (arrows denote drug administration). Plasma levels were measured at 18 and 42 hr. Samples performed in duplicate with individual values displayed. (**C**) Three 35-day-old fetuses were retrieved from the sow in panel B. Liver homogenates were measured after acetonitrile extraction. Samples were run in duplicate and average values are displayed. The column indicates average of three fetuses +/- SEM.
